# Quantifying relationships between chromosome organization and chromosomal aberrations

**DOI:** 10.1101/223859

**Authors:** Y.A. Eidelman, S.V. Slanina, S.G. Andreev

**Affiliations:** Institute of Biochemical Physics, Russian Academy of Sciences, Moscow, Russia; National Research Nuclear University MEPhI, Moscow, Russia

## Abstract

The question of to what extent large scale structure of interphase chromosomes contributes to chromosomal exchange aberrations is discussed for a long time but still remains unclear. We designed the polymer model of 3D organization of a mouse chromosome and simulated X-ray induced chromosomal aberrations exploring two alternative hypotheses: the probability of contacted damaged loci entailing a chromosomal rearrangement is (a) a constant value or (b) not constant, depending on the distribution of DNAsel-hypersensitivity peaks along the chromosome. The latter hypothesis proved to explain the experimental data better than the former, meaning that not only large-scale structure but also local chromatin alterations play a role.

## INTRODUCTION

The relationship between spatial structural organization of chromosomes and radiation-induced chromosomal aberrations (CAs) has long been discussed in the literature, e.g. [1–3]. A comparison between Hi-C data on intrachromosomal contacts and the data on the distribution of radiation-induced breakpoints has shown that they do correlate, but this correlation is not too high [4]. Thus, on one hand, locus contacts (and, consequently, the large-scale three-dimensional structure) are important contributor to CA formation [2], but, on the other hand, there are supposedly other factors. This paper deals with the problem of identifying these possible factors. We developed a computer 3D model of mouse chromosome 2 and verified it according to the experimental data. Then, using the mechanistic model of radiation-induced CAs developed previously [5, 6], we calculated the distribution of the breakpoints for intrachromosomal exchange aberrations based on two hypotheses: a) the probability that the contact of two damaged loci will lead to formation of an exchange aberration is a constant value; b) this probability depends on the number of DNAsel-hypersensitivity peaks. The latter hypothesis proved to explain the experimental data better than the former.

## METHODS

### Interphase chromosome structure

The interphase structure of mouse chromosome 2 was modeled using the molecular dynamics (package OpenMM). A chromosome was considered as a polymer chain of elements with 1 Mbp DNA content. The interaction potential between any pair of elements (i, j) consists of two components, U_ij_(r) = U_ev_(r) + U_attr_(r, i, j), where U_ev_(r) is an excluded volume potential, U_attr_(r, i, j) is an attraction potential depending on the position of the elements along the chain. The excluded volume between any two elements was modeled by the shifted Lennard-Jones potential. The attraction potential was modeled by the attracting component of the Lennard-Jones potential with a coefficient depending on the position of the elements along the chain. To determine these coefficients, we developed an iterative algorithm consisting of the following steps:

1. Initial coefficients are equal for all element pairs.
2. The ensemble of 500–2000 conformations is simulated.
3. The contact map is calculated and compared with the experimental one.
4. If Pearson’s correlation between the contact maps R>0.98, the simulation is stopped. Otherwise, for each element pair (i, j) the coefficient in the attraction potential increases if the simulated contact frequency is lower than the experimental one and decreases if it is higher.
5. Return to step 2.

### Radiation-induced chromosomal aberrations

We used the previously developed [5, 6] algorithm of chromosomal exchange aberration (CA) modeling taking spatial organization of interphase chromosomes into account. Each simulation was performed for 25 million cells (50 million chromosomes 2) which was close to the number of cells in the experiment [4]. As the experimental distribution of intrachromosomal breakpoints (181, j) (i.e. between the element located 181 Mbp apart from p-telomere and any other element j) was built with 2 Mbp resolution, the theoretical distribution was also integrated to this resolution.

## RESULTS AND DISCUSSION

### Interphase chromosome structure

The MD model of the three-dimensional structure of mouse chromosome 2 produces contact maps (i.e., tables with frequencies of contacts between all pairs of loci (i, j)) quantitatively consistent with the experimental data (Fig. 2A, B). Also, the good agreement is demonstrated for the distribution of the contact frequency (181, j), Fig. 1C. The terms used hereafter: “locus 181” is the 1 Mbp region located 180-181 Mbp apart from p-telomere; “frequency (181, j)” is frequency of contacts between locus 181 and other locus with position j (Mbp) along the chromosome. The distribution of the contact frequency (181, j) is of interest to us, as this is the locus for which the experimental data on the breakpoints [4] have been obtained. The possible explanation for higher frequency of contacts for j close to 181 could be local condensation near q-telomere of chromosome 2, or higher probability to come into contact for two elements with low genomic separation compared to that for high genomic separation, or both. The clarification of detailed reasons is the subject for further consideration.

**Fig. 1.**
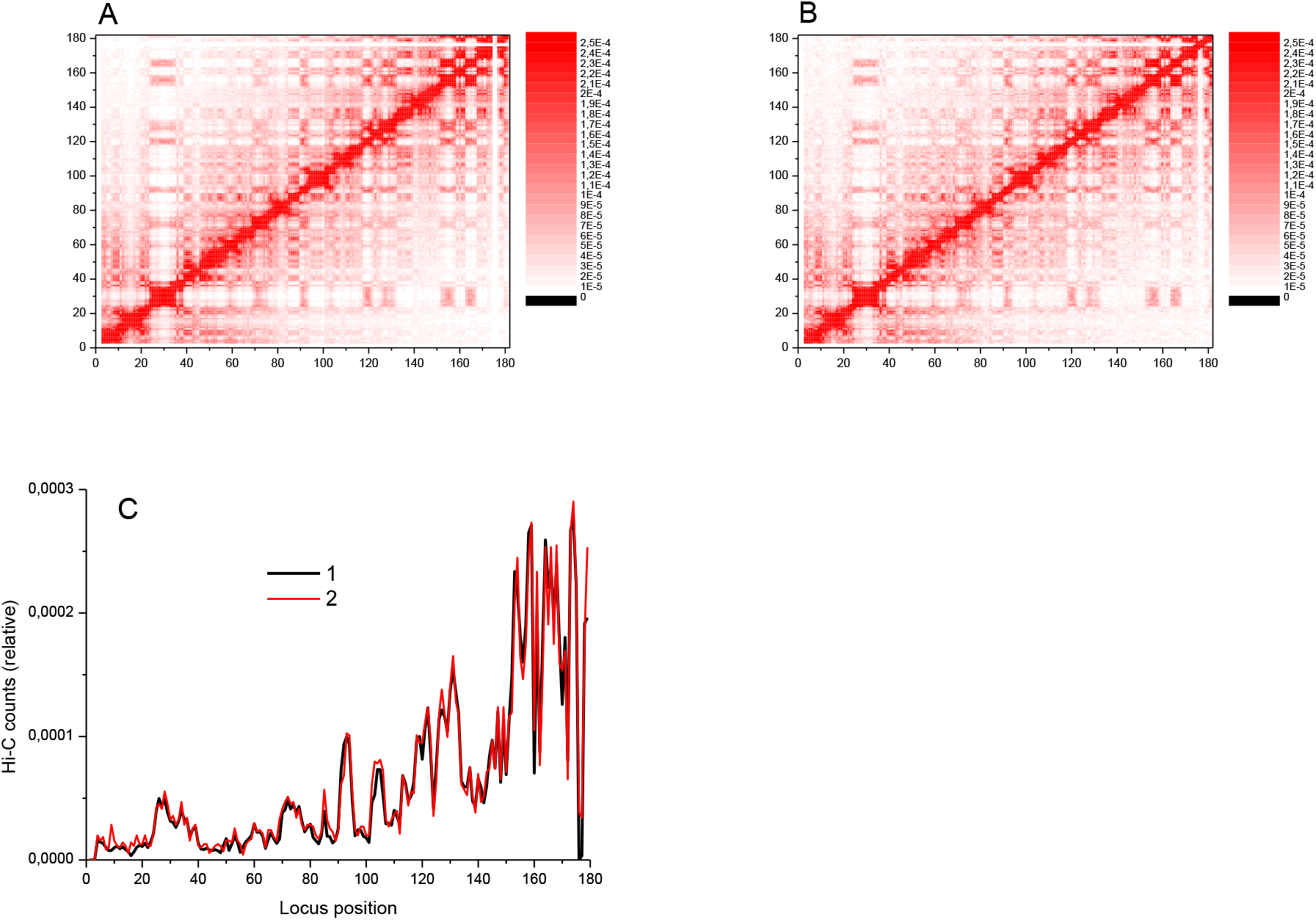
The polymer model of mouse chromosome 2 is consistent with the experimental Hi-C data. A: the contact map for mouse chromosome 2 obtained in Hi-C experiment [4]. Map resolution is 1 Mbp. The map is normalized to the total number of intrachromosomal contacts except those between loci located less than 2 Mbp apart. B: the contact map obtained from an ensemble of 2000 structures simulated, the same normalization as in A. Pearson correlation R=0.982. C: contact frequency between element 181 and other elements as a function of element position along chromosome 2. The same normalization as in A. 1 – experiment [4], 2 – simulation. Pearson correlation R=0.987.

**Fig. 2.**
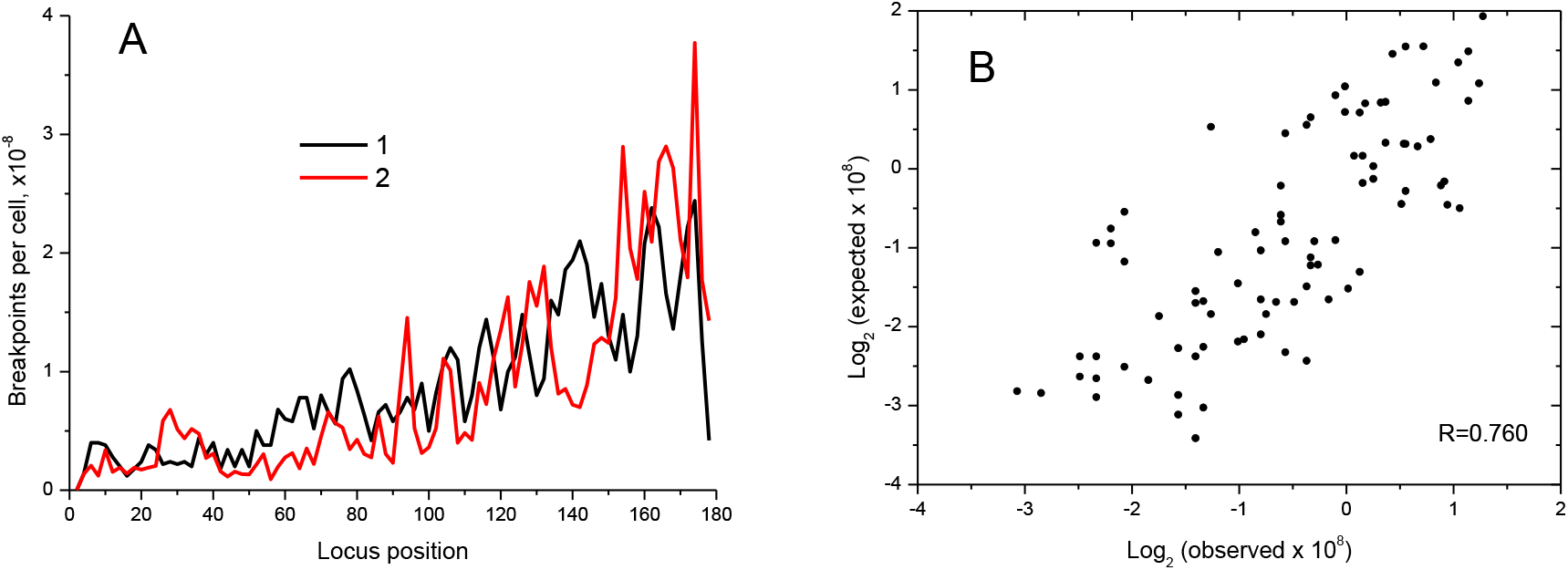
Comparison of the calculated and experimental breakpoint distributions on the basis of the hypothesis of a constant contact-exchange probability. A: the distributions of the breakpoints (181, j). 1 - experiment [4], 2 - simulation. B: the correlation graph between simulation and experiment. Pearson correlation R=0.760.

### Radiation-induced chromosomal aberrations

Distributions of intrachromosomal breakpoints with the participation of locus 181 of chromosome 2, induced by gamma ray exposure with dose 5 Gy, were calculated on the basis of the simplest assumption that the probability of contact-exchange is a constant, i.e. any two contacting loci that contain DNA DSBs can interact to form an exchange aberration with the same probability. The shape of the calculated breakpoint distribution (Fig. 2A) repeats the shape of the experimental distribution of contacts (Fig. 1C), but it does not agree well with the experimental distribution of the breakpoints, see Fig. 2.

To search for possible factors that govern non-constant probability contact-exchange, we plotted the ratio of the experimental number of breakpoints (181, j) and Hi-C counts (181, j) as a function of j (Fig. 3A, curve 1). For breakpoint calculations, “locus j” is referred to as a 2 Mbp block within the chromosome, positioned j Mbp counted from p-telomere. We used 2-Mbp block size because we took the breakpoint distribution with this resolution from [4]. The ratio breakpoints-to-counts was deemed to be the probability contact-exchange. This hypothesis is valid if aberration formation is contributed mainly by contacts of chromosomal lesions immediately after irradiation (“contact-first” mechanism). The dynamical aspects of aberration formation (“breakage-first” mechanism) were not considered in this work.

**Fig. 3.**
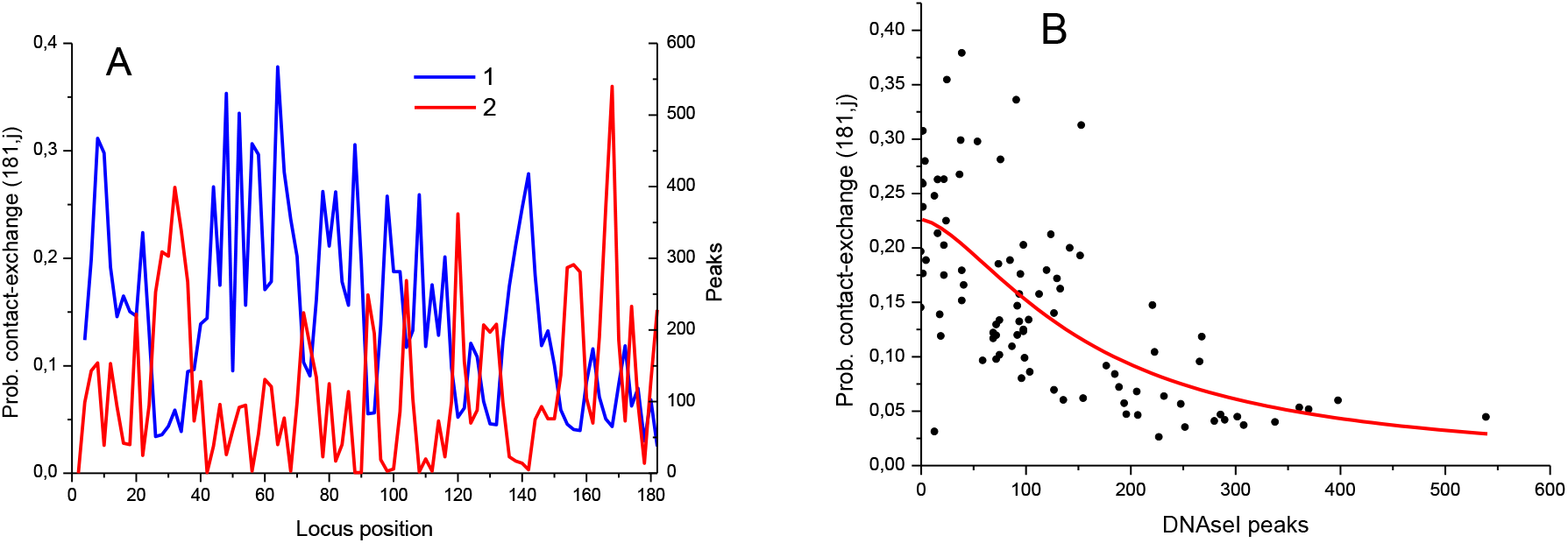
The relationship between chromosome-wide distributions of contact-exchange probability P(181,j) and the DNAsel-hypersensitivity peaks. A: 1 – contact-exchange probability (181, j) *vs* locus position j, data from [4], 2 – total number of peaks of DNAsel-hypersensitivity peaks per 2Mb-locus as a function of locus position along the chromosome. Both graphs are built with 2 Mbp resolution. B: the correlation graph between contact-exchange probability and the number of DNAsel-hypersensitivity peaks. Pearson correlation R = −0.648. The red curve is the approximating function (1).

We checked how the ratio breakpoints-to-counts, or the probability contact-exchange (181, j), correlates with distribution of DNAseI-hypersensitivity (HS) peaks. It is often observed that chromatin containing many HS-peaks is in more open state and more transctiptionally active than chromatin containing few HS-peaks (e.g. [7]). The information on HS-peak positions, available with high resolution (~10 kbp), was taken from ENCODE [8], and then number of peaks was summated over each 2 Mbp block. The ENCODE data were taken for chromosome 2 of mouse spleen B cells since the HS-data on A-MuLV-transformed ATM^-/-^ pro-B cells used in [4] were unavailable.

The distribution of HS-peaks number in 2 Mbp blocks (loci) *vs* locus position along chromosome 2 is shown in Fig. 3A (curve 2). There was an anticorrelation between the probability contact-exchange (181, j) and the number of peaks in locus j, R = −0.648 (Fig 3B). Thus, by this criterion the more active locus j is, the less aberrations are formed between the pair (181, j).

From the correlation graph (Fig. 3B) we approximated the dependence of the contact-exchange probability on the number of peaks by the function

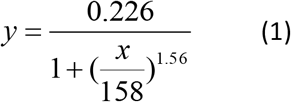

On the basis of this dependence, we recalculated the distribution of the breakpoints (181, j) and compared it with the experimental one, see Fig. 4.

**Fig. 4.**
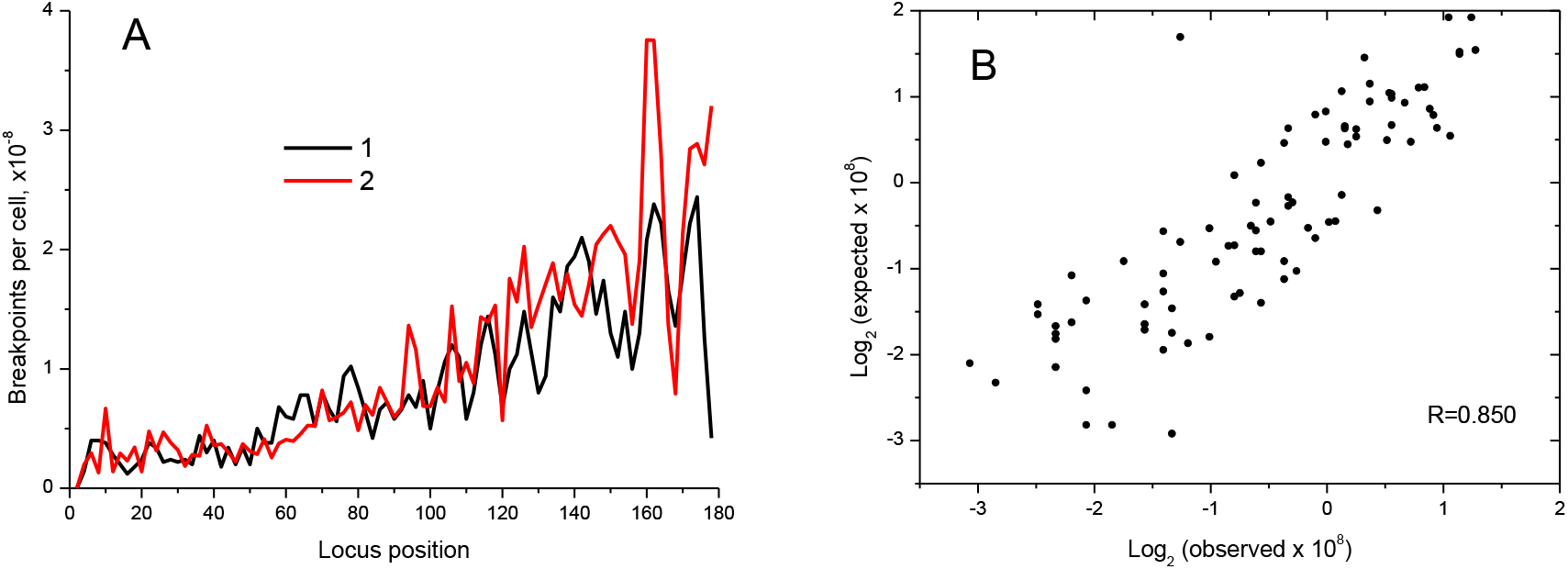
Comparison between calculated and experimental breakpoint distributions on the basis of the hypothesis of contact-exchange probability nonhomogeneous, depending on the distribution of DNAsel-hypersensitivity sites. A: the distribution of the breakpoints (181, j) along the chromosome (locus position). 1 - experiment [4], 2 - simulation. B: the correlation graph between simulation and experiment. Pearson correlation R=0.850.

Fig. 4 demonstrates that the hypothesis of an inverse dependence of the contact-exchange probability on the number of hypersensitivity peaks leads to much better agreement with the experimental data in comparison with the initial hypothesis of the contact-exchange probability being constant, the Pearson correlation coefficient R = 0.850 vs 0.760. However, in some areas the discrepancies are still significant. This suggests that, in addition to local chromatin alterations, some other factors play a role which are yet to be identified.

## ACKNOWLEDGEMENTS

The present work was supported by Russian Foundation for Basic Research grant 14-01-00825 to S.A. S.A. acknowledges support from the MEPhI Academic Excellence Project (Contract No. 02.a03.21.0005).

